# In Silico Design of Self-Optimizing Integrated Metabolic and Gene Regulatory Networks

**DOI:** 10.1101/007658

**Authors:** Timo R. Maarleveld, Bennett K. Ng, Herbert M. Sauro, Kyung Hyuk Kim

**Author notes:** Corresponding author Email: Timo R. Maarleveld* –; Bennett K. Ng –; Herbert M. Sauro –; Kyung Hyuk Kim –.

## Abstract

Biological organisms acclimatize to varying environmental conditions via active self-regulation of internal gene regulatory networks, metabolic networks, and protein signaling networks. While much work has been done to elucidate the topologies of individual networks in isolation, understanding of inter-network regulatory mechanisms remains limited. This shortcoming is of particular relevance to synthetic biology. Synthetic biological circuits tend to lose their engineered functionality over generational time, primarily due to the deleterious stress that they exert on their host organisms. To reduce this stress (and thus minimize loss of functionality) synthetic circuits must be sensitive to the health of the host organism. Development of integrated regulatory systems is therefore essential to robust synthetic biological systems. The aim of this study was to develop integrated gene-regulatory and metabolic networks which self-optimize in response to varying environmental conditions. We performed *in silico* evolution to develop such networks using a two-step approach: (1) We optimized metabolic networks given a constrained amount of available enzyme. Here, we found that a proportional relationship between flux control coefficients and enzyme mass holds in all linear sub-networks of branched networks, except those sub-networks which contain allosteric regulators. Network optimization was performed by iteratively redistributing enzyme until flux through the network was maximized. Optimization was performed for a range of boundary metabolite conditions to develop a profile of optimal enzyme distributions as a function of environmental conditions. (2) We generated and evolved randomized gene regulatory networks to modulate the enzymes of a target metabolic pathway. The objective of the gene regulatory networks was to produce the optimal distribution of metabolic network enzymes given specific boundary metabolite conditions of the target network. Competitive evolutionary algorithms were applied to optimize the specific structures and kinetic parameters of the gene regulatory networks. With this method, we demonstrate the possibility of algorithmic development of integrated adaptive gene and metabolic regulatory networks which dynamically self-optimize in response to changing environmental conditions.

## Introduction

In synthetic biology living organisms have been engineered to perform novel functions that do not exist in nature. Some synthetic constructs include tunable oscillators [1], event counters [2], concentration detectors [3], edge detectors [4], linear amplifiers [5], and toggle-switches [6]. Synthetic genetic circuits, however, tend to lose their engineered functionality over time, due in large part to their deleterious effects on host fitness [7, 8]. Wild-type cells and mutant synthetic cells often out-compete their designed synthetic counterparts. One possible method to improve this fitness stability is to optimize the expression of synthetic gene products to reduce the burden on the host cell. For this, the interaction between the synthetic circuits and host metabolic networks must be considered.

Most micro-organisms live in continuously changing environments, which makes maintaining fitness even more difficult. An example of this are the phototrophic micro-organisms cyanobacteria that are subject to the daily fluctuations in environmental light intensity. Recently, cyanobacteria have been engineered to convert carbon dioxide into biofuels with designed synthetic pathways [9, 10]. For efficient biofuel production, the synthetic pathways must be adaptive to this varying environmental condition. The adaptive responses, controlled by gene regulatory networks (GRNs), allow the host to quickly shift from one gene expression profile to another as needed. For instance, upon a shift from light to dark, the cyanobacterium *Synechococcus* sp. PCC 7002 responds by decreasing and increasing the transcription levels of genes essential to photosynthesis and carbohydrate degradation [11]. To understand the response mechanisms of such adaptive systems, it is important to adopt an integrative perspective. The nature of GRNs, protein signaling networks, and metabolic networks must be appreciated as an interconnected whole. In this study we adopt this perspective by considering GRNs within an integrated metabolic context.

The linkage between GRNs and their target metabolic networks are catalytic enzymes. Accurate expression of these catalytic enzymes for a wide range of environmental conditions gives robustness to the micro-organism. The concentrations of these enzymes are thought to be optimized for efficient cellular functions [12, 13]. Optimization is required since protein synthesis and degradation are energetically costly and resource-limited. The availability of amino acids, within the finite volume of a cell constrains the total enzyme concentration within the cell [14]. Additionally, the production of certain osmotically-active enzymes must be constrained to preserve the osmotic balance of the cell [15, 16].

Extensive research has been performed to examine the effect of enzyme mass constraints on metabolic fluxes. Several different optimization conditions have been considered in those studies: Metabolic fluxes were maximized for a fixed total enzyme concentration [17, 18]; intermediate concentrations and total osmolarity were minimized [16]; net enzyme concentrations were minimized [12, 13] or maintained below a certain level [14]. The studies which sought minimal total enzyme concentration as an optimization condition [12,13] show clear relationships between enzyme concentrations and flux control coefficients, namely, that flux control coefficient of any enzyme is proportional to that enzyme’s concentration in the optimized state. It was shown, however, that this relationship is valid for linear pathways but not for branched pathways. Consequently a new relationship was derived for general network topologies [13].

We aimed to design GRNs to reduce the burden a synthetic circuit exerts on the host cell, and to make synthetic systems robust to a wide range of environmental conditions. To achieve this aim, we considered the interaction between GRNs and metabolic networks, and investigated the structural properties of designed GRNs. These GRNs were designed to regulate the expression of catalytic enzymes such that metabolic fluxes remained maximized for various boundary metabolite concentrations. By taking an *in silico* evolutionary approach we obtained integrated networks composed of a GRN and a metabolic network which are robust and self-optimizing (Figure 1). We found that the integrated networks have interesting topological properties: the GRN typically shows shallow regulation and feed-forward structures, and boundary metabolites alone can be sufficient for regulation of the GRN.

**Figure 1:**
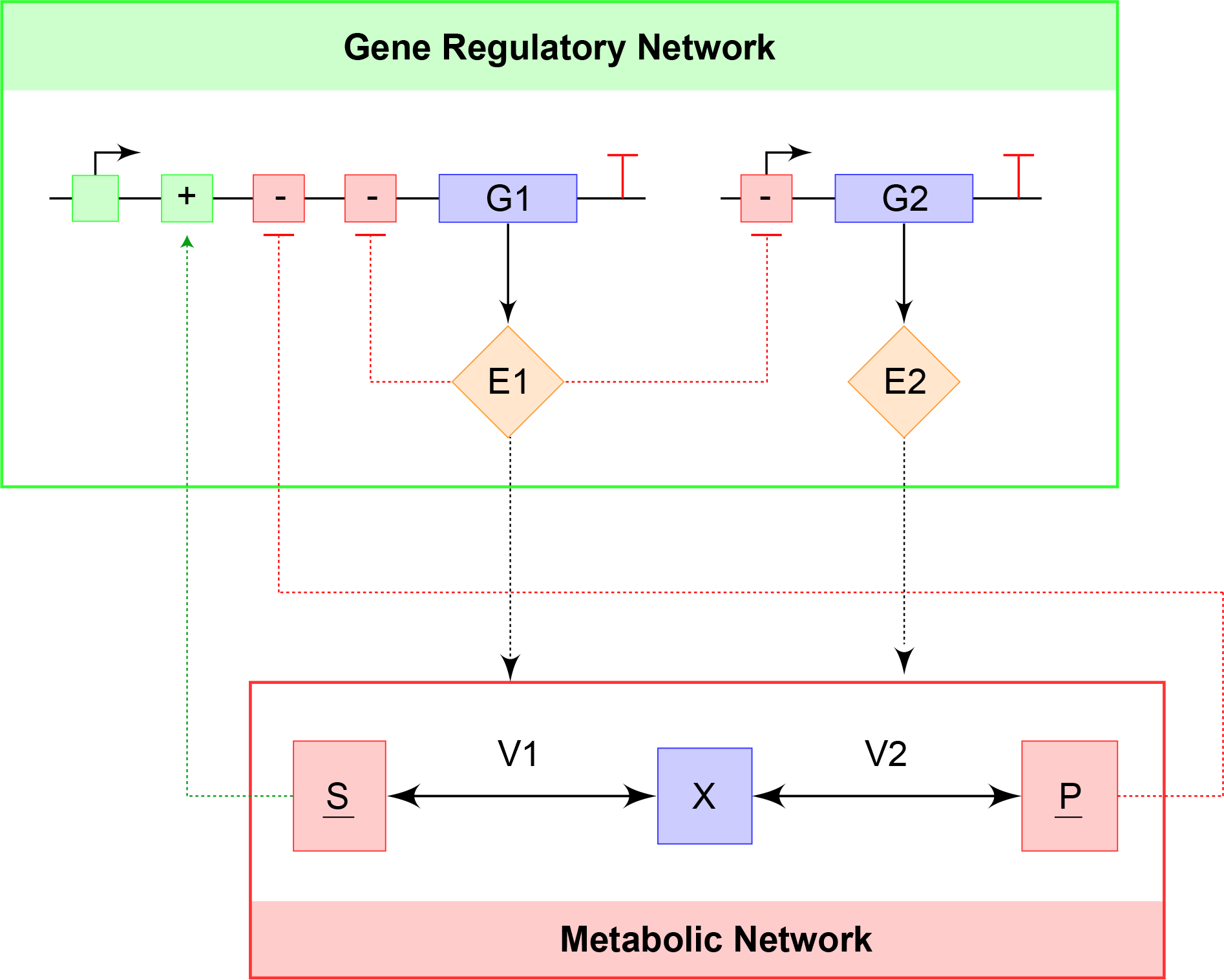
Self-optimizing integrated GRN-metabolic networks: The GRN regulates the expression levels of both enzymes (*E*_1_ and *E*_2_) such that the enzyme distribution maximizes the metabolic flux for a given total enzyme mass. This regulation is performed by monitoring the metabolic network. In this study, we consider the boundary metabolites as input signals for the GRN. Floating and fixed metabolites are indicated in blue and red squares, respectively. Rate equations *v*_1_ and *v*_2_ are described by reversible Michaelis-Menten kinetics, where *E*_1_ and *E*_2_ catalyze reactions *v*_1_ and *v*_2_, respectively.

Moreover, our numerical study discovered a new relationship similar to the one found in the linear pathway study [12, 13] which holds for general networks that include both linear and branched pathways. This relationship was found to be valid for each branch of a network which meets two conditions: (1) the branch itself is linear and (2) the branch does not include a metabolite which that allosterically regulates another branch. By investigating local linear pathways of a general network, this relationship can be used to determine whether pathway metabolites regulate other parts of the network.

## Results

### Model systems

A self-optimizing network that maximizes metabolic fluxes under a constraint in the total enzyme mass is composed of two reaction systems: a GRN and a metabolic network (Figure 1). The GRN controls the expression of enzymes that catalyze metabolic reactions and the metabolic network regulates promoter activities in the GRN via allosteric metabolite interactions. The metabolic and gene regulatory networks were described in a deterministic framework which does not consider the stochastic details of gene expression [19]. The networks were modeled with a set of ordinary differential equations with non-linear reaction rates (see Methods for details).

### Flux control coefficients reflect enzyme concentrations

Previous studies which considered the total enzyme mass constraint revealed a relationship between flux control coefficients and enzyme concentrations in *linear* pathways. Brown [12] proposed that flux control coefficients [20–22] become proportional to enzyme concentrations in the optimized state for an n-step *linear* metabolic pathway:

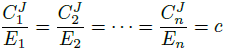

where *c* is a constant and every enzyme has identical mass. The flux control coefficient 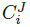 describes the sensitivity of the steady-state flux *J* to a perturbation in the concentration of the *i*-th enzyme *E_i_*:

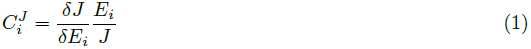

The control coefficient has been used to quantify the degree of control specific reactions exert on fluxes within a network [23], as well as control of gene expression noise in gene regulatory networks [24, 25].

The above relationship can be extended to consider enzymes of unequal mass as follows:

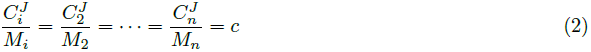

Where *M_i_* = *E_i_m_i_* with *m_i_* the molar mass of the *i*-th enzyme. In the previous study [12], total enzyme mass was minimized for a given metabolic flux. Equation (S14) can also be derived for the complementary case where flux is maximized for a given total enzyme mass (*SI, Section S1*). For *general* metabolic networks, Klipp and Heinrich [13] found a similar relationship which can also be expressed with unequal enzyme mass (*SI, Section S1*):

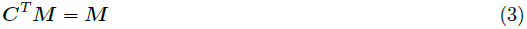

where ***C*** represents a matrix of flux control coefficients 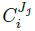 and ***M*** represents a vector of enzyme masses *M_i_*.

We investigated the above results numerically and found an interesting fact: Equation (S14) holds for certain sub-networks of general networks which include linear and branched pathways. This previously-unobserved result is a hidden property of Equation (S15). First, consider the branched network shown in Figure 2(e) without allosteric regulation. We minimized the total enzyme concentration while all the fluxes were fixed (see Methods for details). The optimization procedure yielded Brown’s result for all branches (Figure 3(a)). This can be partly understood with the following intuitive argument: When the total mass of enzymes regulating each branch is minimized, the total mass of enzymes for the whole network must also be minimized. Since each branch flux is fixed, the optimization produces Brown’s result for each branch.

**Figure 2:**
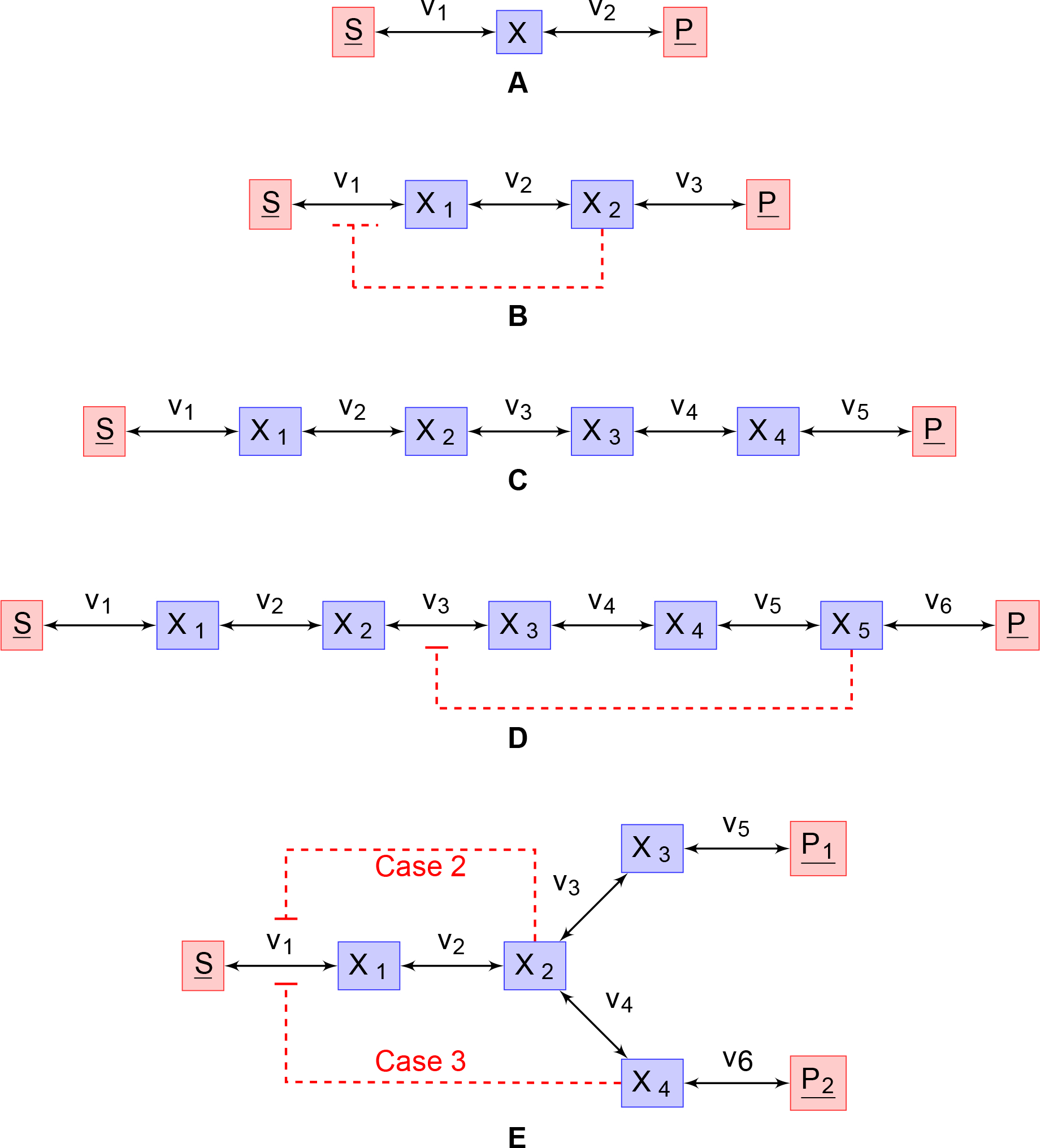
Input metabolic networks. (a) A two-reaction metabolic pathway regulated by two enzymes (*E*_1_ and *E*_2_). *S* and *P* are fixed metabolites (red-boxed), and *X* is a floating metabolite (blue box). (b-d) A three-reaction metabolic pathway with negative feed-back, a five-reaction metabolic pathway, and a six-reaction metabolic pathway with negative feed-back were considered. (e) A branched network was explored for three different cases: (Case 1) Without feed-back, (Case 2) with negative feed-back from *X*_2_ on *V*_1_, and (Case 3) with negative feed-back from *X*_4_ on *V*_1_

**Figure 3:**
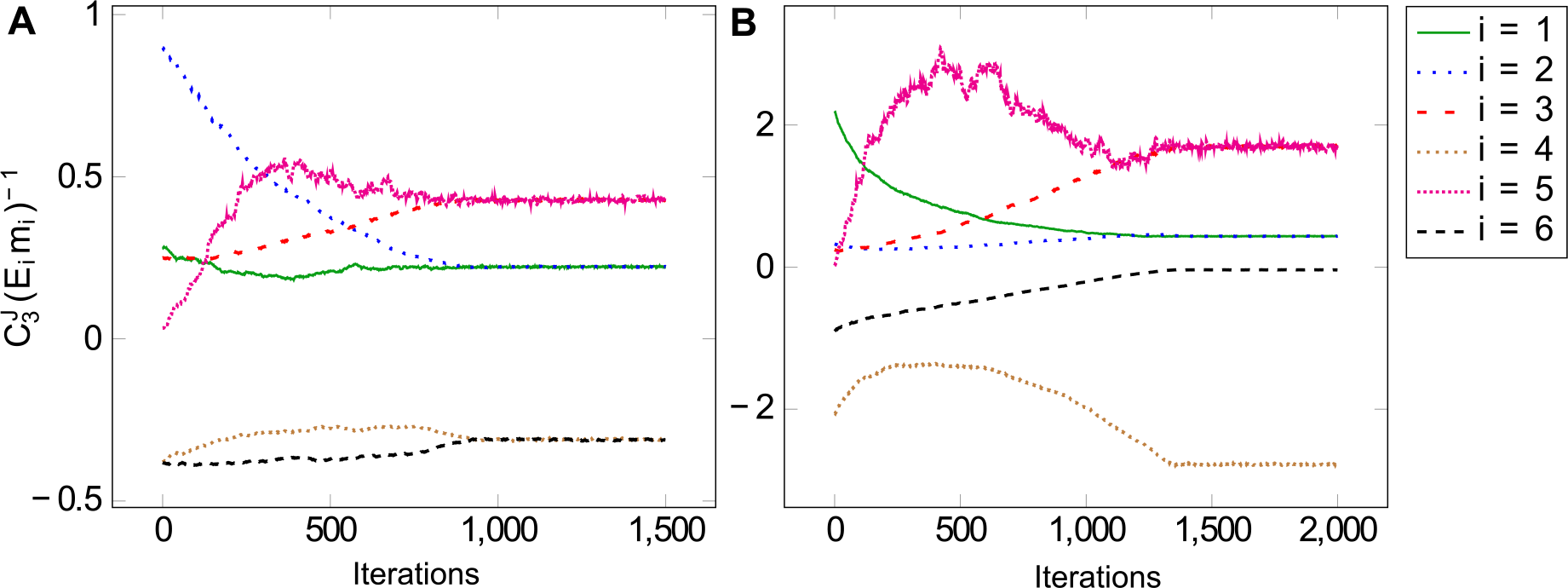
Proportionality between flux control coefficients and enzyme concentrations in the **branched network shown in** Figure 2(e). (a) Without allosteric regulation (Case 1 in Figure 2(e)): Brown’s result holds for all linear sub-networks. (b) With allosteric regulation (Case 3): Brown’s result does not hold from the branch were the regulation originates. See Figures S3 and S5 for details of all flux control coefficients.

For the branched network shown in Figure 2(e), we examined various different regulatory configurations (Figure 3; Figures S3 – S5). Surprisingly, we observed the same result for all branches which are the target of inter-branch allosteric regulation, but not for those branches which are the source of allosteric regulation. Brown’s result also holds for all branches if a branch point corresponds to an allosteric metabolite (Figure S4). Mathematically, 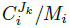 were equal for all metabolites within any linear segments of the networks that do not have any allosteric metabolite regulating other branches. We also numerically solved Equation (S15) and verified our simulation results (*SI*).

These findings suggest that our numerical relationship can be used to determine whether metabolic networks are in an optimized state by observing the control coefficients of each linear branch, excluding those with allosterically-active metabolites. In addition, this relationship can be leveraged to discover the origins of regulation (allosteric metabolites) within a network., under the condition that total enzyme mass is tightly constrained.

The connectivity theorem [20–22] suggests that reactions with low enzyme levels have correspondingly high flux control coefficients. But our results and previous studies [12, 13] show that reactions with higher control coefficients have higher enzyme levels. At first, this seems paradoxical. Consider the two-reaction linear pathway shown in Figure 2(a). We computed the control coefficients for each reaction over a range of different enzyme concentrations, where the net enzyme mass was fixed. Figures 4(a–b) illustrate that higher enzyme levels can lead to lower flux control coefficients.

**Figure 4:**
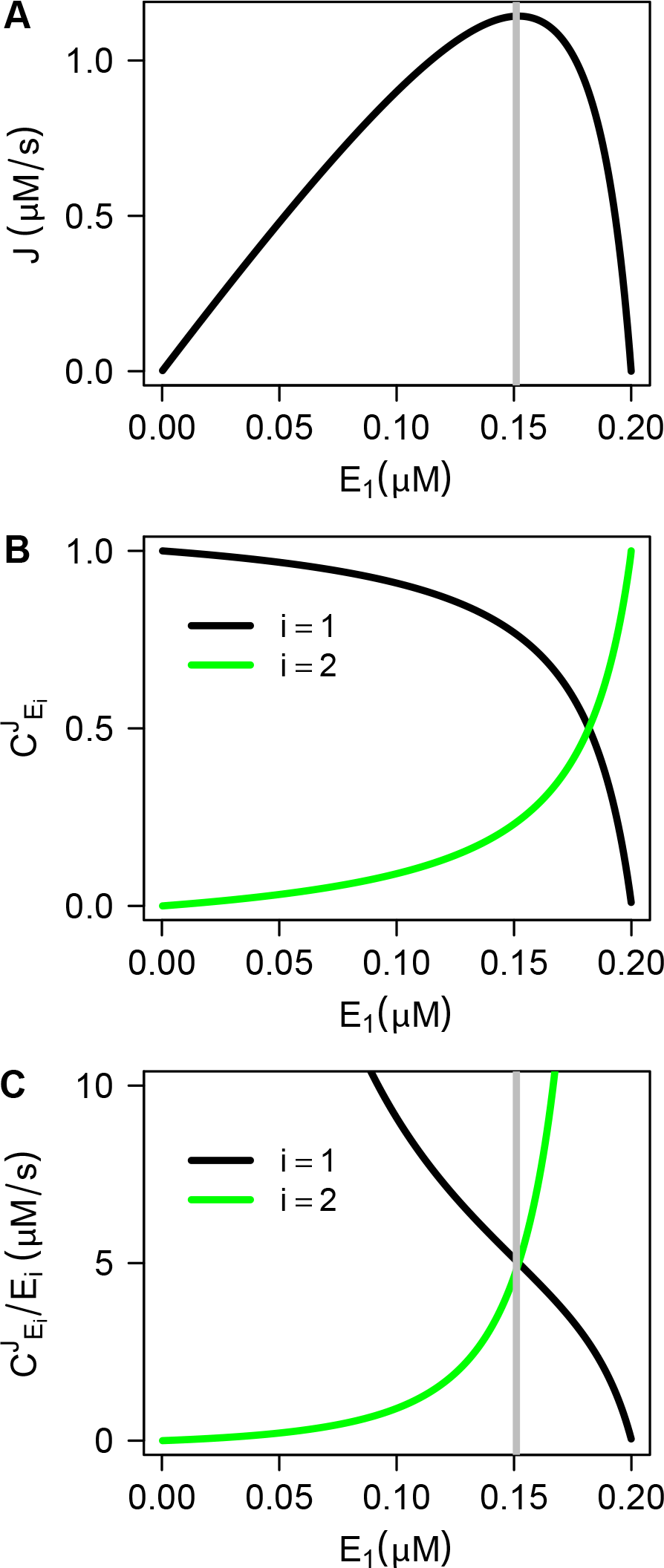
Control coefficients at the flux maximum: (a) In a two-reaction metabolic pathway shown in Figure 2(a), the flux became maximized under the total enzyme mass constraint. Here *E*_1_ and *E*_2_ were assumed to have identical mass. (b) As enzyme concentration increases, its corresponding flux control coefficient decreases for each *E*_1_ and *E*_2_. (c) At the maximum flux level, 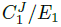 matched 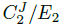.

The flux peaks at an intermediate *E*_1_ concentration (Figure 4(a)). The flux control coefficient for *E*_1_, 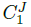 decreases monotonically with the increase in *E*_1_ (Figure 4(b)). This illustrates that higher enzyme levels can lead to lower flux control coefficients of that particular enzyme. However, this relationship between an enzyme level and its corresponding flux control coefficient can only be observed with changes in the enzyme concentration, not with a concentration value in isolation. Equation (S14), on the other hand, describes the latter case. Consequently, the Brown’s result is biologically reasonable.

We have hitherto investigated the properties of flux control coefficients under the total enzyme mass constraint while optimizing metabolic fluxes. In the following sections, we will investigate how such an optimized state can be maintained under a wide range of environmental conditions.

### In silico evolution of GRNs as a part of integrated self-optimizing networks

We composed gene regulatory networks and metabolic networks to obtain integrated networks which function robustly under a wide range of environmental conditions. We optimized each metabolic network over a wide range of boundary metabolite concentrations to determine corresponding optimal enzyme distributions (see *SI, Section S5*). These data were used then as inputs and outputs for *in silico* evolution of GRNs [26, 27]. Here the boundary metabolite concentration and total enzyme mass are considered environmental conditions.

*In silico* evolution of GRNs was performed by building *N* random GRNs from a given limited number of gene network building blocks and parameterizations. The building blocks were (1) genes that express enzymes, (2) inducible, repressible, and constitutive promoters, and (3) up to three additional regulator binding sites. An example of such a random GRN is shown in Figure 1.

Given a specific boundary metabolite condition, the constructed GRNs were trained to produce the corresponding optimal enzyme distribution. The fitness of each GRN was quantified as the relative error between the desired and trained output values, as expressed by Equation (4). After evaluating network fitness, some of the networks were selected to breed for a new generation (Figure 8(b)). Fitter networks were more likely to survive and mutate into even fitter GRN structures. Two selection methods — 1) elitism to guarantee survival of the fittest structures and 2) tournament selection to allow structures to mutate to fitter structures — were used to select GRNs for the next generation. The GRNs chosen with tournament selection were subject to mutations in parameter values and regulation patterns. Activators could be mutated into repressors (and vice versa), and regulators or genes could be inserted or deleted within predefined bounds. Typically, the numbers of genes and regulators allowed were two and three times of the number of enzymes regulating the connected metabolic networks, respectively. The process of fitness-scoring, network selection, and reproduction was iterated until the fitness score converged. Finally, the gene and metabolic network were integrated, resulting in a self-optimizing biochemical network. See the Methods for more details on this evolutionary approach.

For the metabolic networks shown in Figure 2(a–d), a GRN was trained to fit the desired input-output characteristics. One of the most surprising findings was that no additional proteins were required in the GRN to satisfy the desired input-output characteristics. For example, to control the input-output characteristics in the two-enzyme metabolic network shown in Figure 2(a), the gene networks were initially allowed to contain up to four genes. Two of these genes would express both the enzymes while other genes could express enzymes that do not interact with the metabolic network. Nonetheless, the evolutionary approach returned a GRN in which two genes were sufficient to maximize the metabolic flux in the trained regime of boundary metabolite concentrations. This result shows that the enzymes expressed in the GRNs can play dual roles of regulation on themselves as well as on the target metabolic networks, which leads to shallow regulation cascade structures.

The input-output performance of the evolved GRNs was evaluated by computing the degree of fitness (performance error; Equation (4)) for a wide range of boundary metabolite concentrations beyond the trained regime. The performance error is shown in Figure 5, where the dotted white boxes represent the trained regimes. Within the trained regime performance error could be lowered below 1% for all evolved GRNs, indicating excellent input-output performance within this regime. The performance errors over the trained regime were 0.57 % (SD = 0.38), 0.85 % (SD = 0.57), 0.51 % (SD = 0.24), and 0.66 % (SD = 0.25) for the four integrated GRN-metabolic networks referenced in Figure 5, respectively. Outside the trained regime (one or two orders of expansion), the error moderately increased up to approximately 20% for most cases.

**Figure 5:**
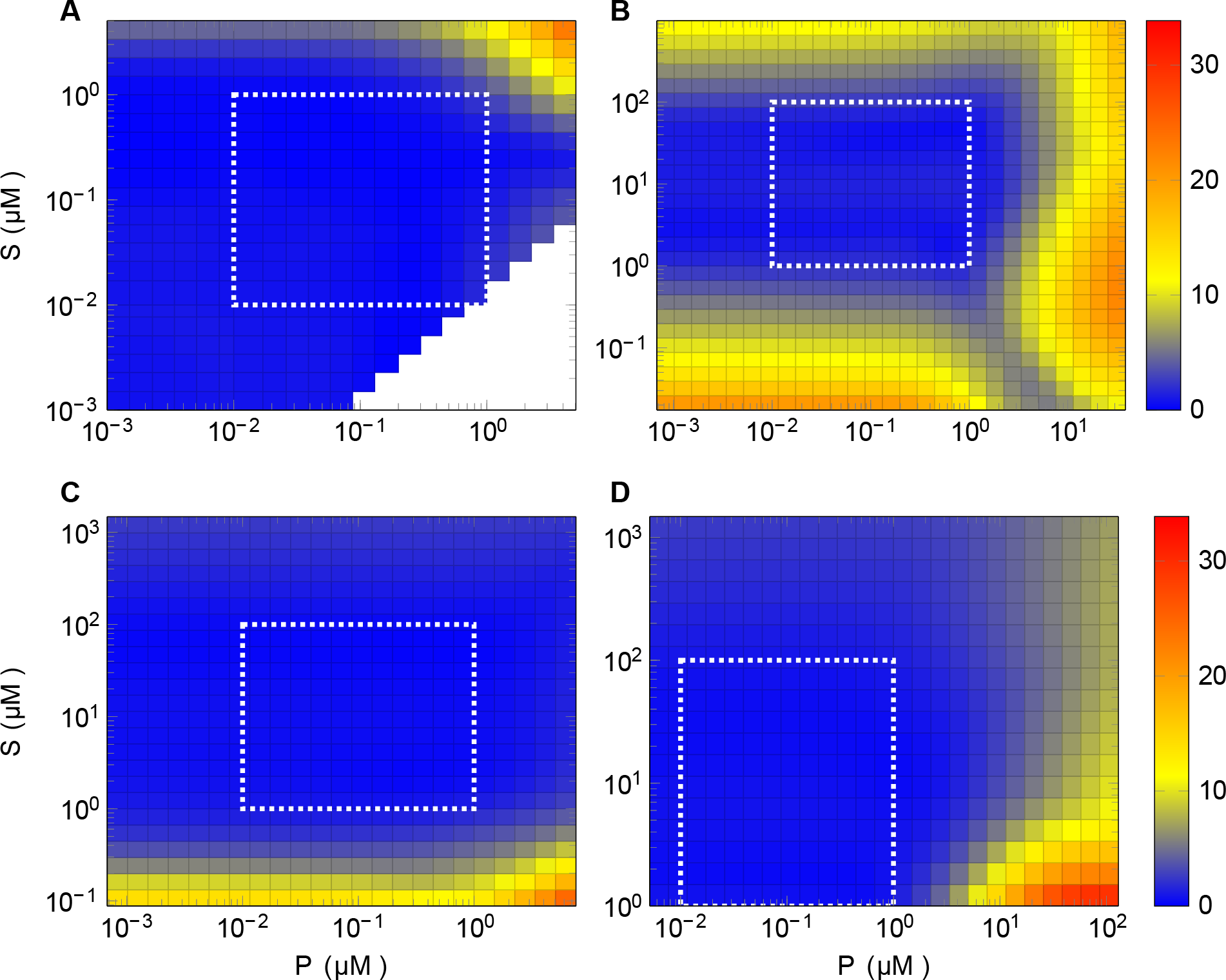
Performance of trained GRNs from the *in silico* GRN optimization approach: Percentage deviation of simulated (trained) enzyme concentrations from their desired (training) values. The boundaries of the trained regime are indicated with dotted white boxes. (a) Two-enzyme GRN shown in Figure 6 (a) was integrated with the metabolic network shown in Figure 2 (a). (b-d) The GRNs described in Figure 7 (a-c) were integrated with metabolic networks with three, five, and six enzymes shown in Figure 2 (b-d), respectively.

### Emerging regulation patterns in integrated self-optimizing networks

Besides seeking strong agreement between desired and trained gene expression levels, we were interested in determining general design features of successfully-evolved GRNs. Such design features are listed below.

1. Regulation by boundary metabolites is sufficient for optimal adaptation. Because the optimal enzyme distribution changes for different boundary metabolite concentrations, direct or indirect regulation by boundary metabolites is essential. Based on our analysis of linear metabolic pathways, GRNs obtained by *in silico* evolution (e.g. Figures 6–7) showed accurate input-output performance for wide ranges of boundary metabolite concentrations. This implies that direct regulation by boundary metabolites alone may be sufficient for optimal adaptation to changes in environmental conditions.
2. Shallow network topologies allow coordinated expression. In the five-enzyme system shown in Figure 7(b), enzyme *E*_5_ regulates the expression of three other enzymes, and in the six-enzyme system shown in Figure 7(c), enzyme *E*_6_ regulates four other enzymes. This emerging regulation motif, previously termed a single-input module (SIM) [28], is notable because it allows for coordinated expression with shallow network topologies. The SIM is also known to lead to fast responses to external signals [29–31]. Although the response speed was not considered in our GRN evolution, this result suggests that the frequent occurrence of the SIM in natural systems may be attributed to their utility for robust self-optimization under various environmental conditions.
3. Feed-forward loops can be essential to obtain optimal enzyme distributions. Feed-forward loops (FFLs) were observed in the evolved networks. For example, Figure 7 (c) shows that boundary metabolite *P* inhibits the expression of *E*_3_, but activates the expression of *E*_6_, which in turn actives *E*_3_, completing a type-3 incoherent FFL. This implies a non-monotonic, bell-shape response profile between *E*_3_ and *E*_6_: As *P* increases, *E*_3_ and *E*_6_ increase together but after a certain threshold in *P*, *E*_3_ begins to decrease due to strong inhibition from *P*, while *E*_6_ still increases due to activation from *P*. The prevalence of incoherent FFLs suggests that this motif is an important component for regulation of complex enzyme distributions.

**Figure 6:**
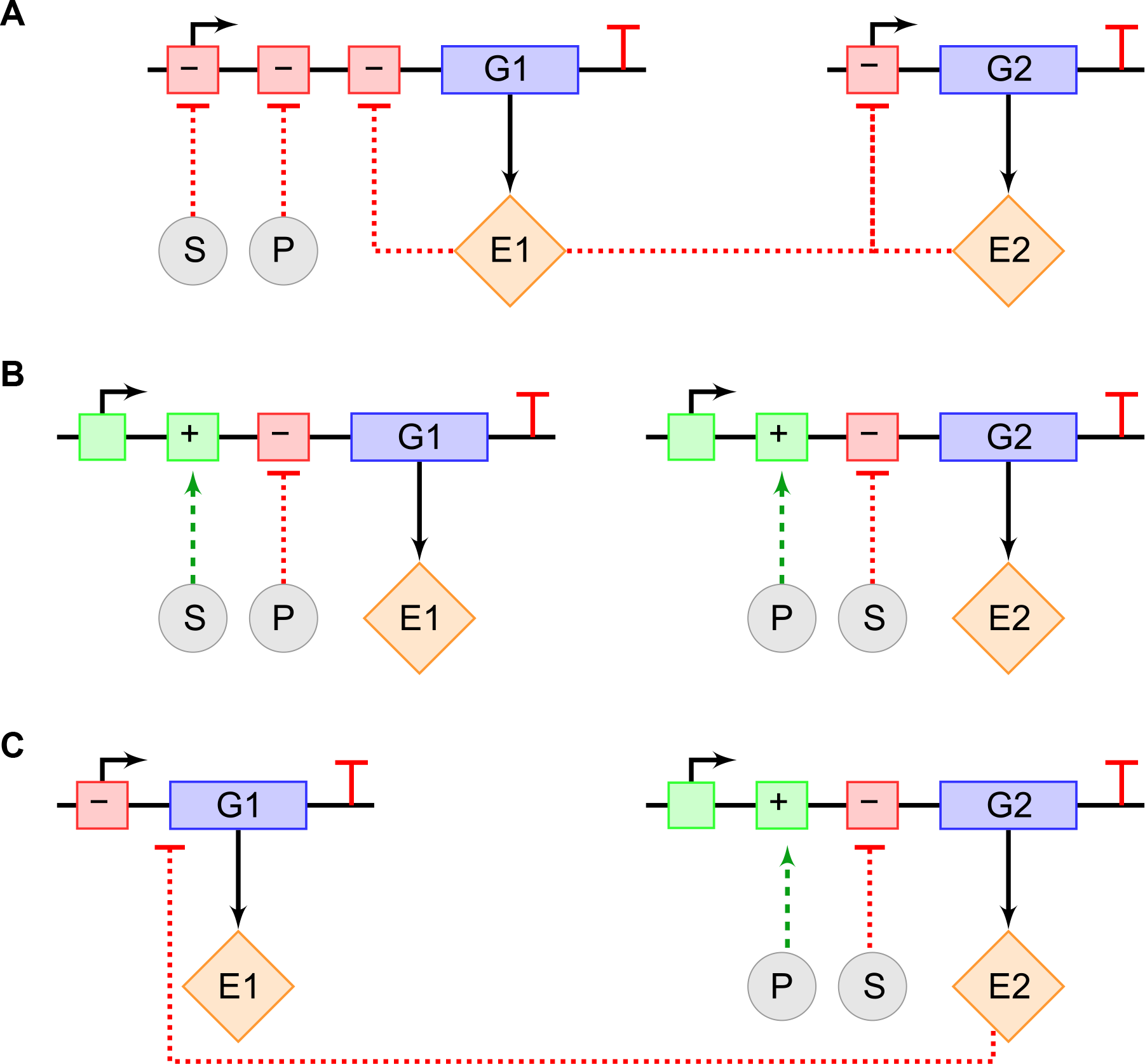
Optimized GRNs capable of maximizing the flux in the two-enzyme metabolic pathway shown in Figure 2(a). Mathematical models are available in the *SI, Section S6*.

**Figure 7:**
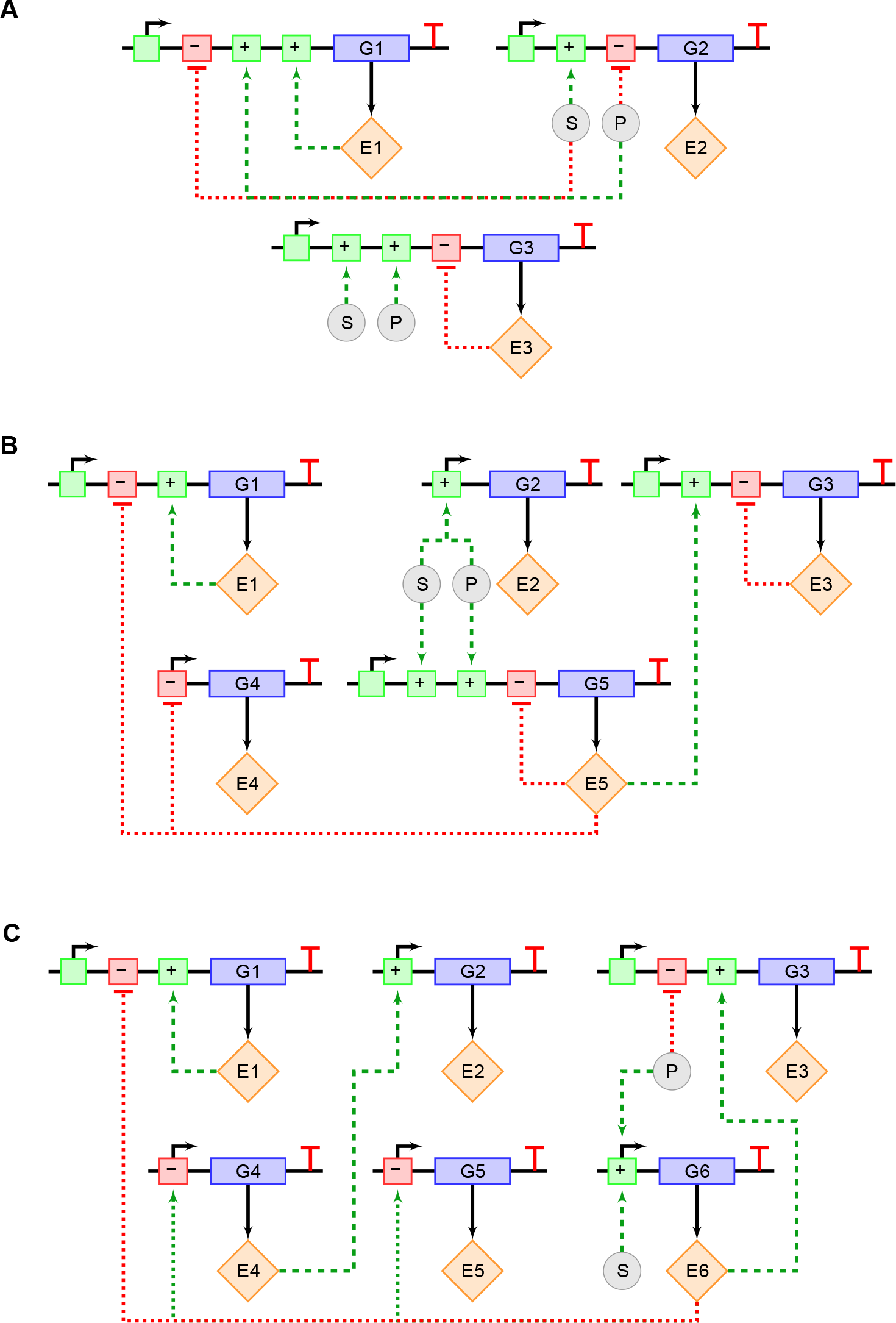
Evolved GRNs for metabolic networks with three or more enzymes. (a) GRN for the three-enzyme metabolic network with negative feed-back (Figure 2(b)); (b) GRN for the five-enzyme metabolic network (Figure 2(c)); (c) GRN for the six-enzyme metabolic network with negative feed-back (Figure 2(d)). Mathematical models are available in the *SI, Section S6*.

## Discussion

This research investigated the design of robust and self-optimizing networks composed of gene regulatory networks and metabolic pathways under the constraint of limited enzyme resources within a cell. The obtained integrated networks adapted to maintain maximal network fluxes for a wide range of boundary metabolite concentrations. Here, boundary metabolite concentrations and the enzyme mass constraint were considered to be directly affected by environmental variations.

Under the total enzyme mass constraint, we computed the optimal enzyme distributions and discovered an interesting relationship between flux control coefficients and enzyme levels for linear sub-networks. The relationship was shown to hold for the sub-networks that (1) are linear and (2) do not allosterically regulate other branches. This relationship can serve as an indicator for the presence of such allosteric metabolites and for determining whether cellular conditions whether are tightly constrained by available enzyme mass.

Furthermore, in our study of self-optimizing integrated networks, we found that a minimal number of metabolites can be used as input signals for self-optimizing GRN controllers of metabolic networks. The boundary metabolite concentrations and corresponding optimal enzyme distributions were used as constraints in evolving GRNs based on genetic algorithms [32]. Here, the initial randomly-built GRNs were evolved *in silico* from a limited number of genetic building blocks and allowable parameterizations. The evolved GRNs were shown to successfully modulate expression levels of the enzymes in response to changes in boundary metabolite concentrations.

One of the important regulation patterns of the evolved GRNs was the SIM, which enabled shallow network regulation structures. This motif implies fast and coordinated regulation. The degree of fitness (performance error; Equation (4)) used in our evolutionary algorithm was related to the steady-state expression levels of the enzymes, and did not include the temporal response to the steady-state. The prevalence of the SIM in the evolved GRNs implies that accurate performance can be obtained by this motif without delaying the dynamic response. It would be interesting to include response time in our fitness function in future research, to investigate emerging GRN motifs related to rapid adaptation of GRNs under volatile environment conditions.

The input-output performance of the *in silico* evolved GRNs was shown to be accurate for wide ranges of boundary metabolite concentrations. The performance plots (Figure 5) show that our GRNs and metabolic networks can be optimized for a range of three or four orders of magnitude within*∼*20% error. This implies that flux optimization can be robustly achieved under significant environmental changes via feedback from GRNs.

In summary, we investigated self-optimizing feedback systems composed of a GRN and a metabolic network under the constraint of limited total enzyme mass. The self-optimizing systems generated accurately maintained optimal enzyme distributions to maximize the metabolic fluxes under significantly different boundary metabolite levels. Our theoretical study showed a possible approach for designing such robust synthetic systems. This work may eventually help provide rational design principles and computational methods for constructing synthetic circuits that are robust to a wide range of environmental conditions.

## Methods

Our self-optimizing integrated GRNs and metabolic networks were designed by using two major algorithms for the optimization of metabolic networks and the *in silico* evolution of GRNs. The first algorithm determined the input-output characteristics of the GRNs and the second algorithm used them as a training data set. The pipeline for designing these self-optimizing networks is shown in Figure 8(a).

**Figure 8:**
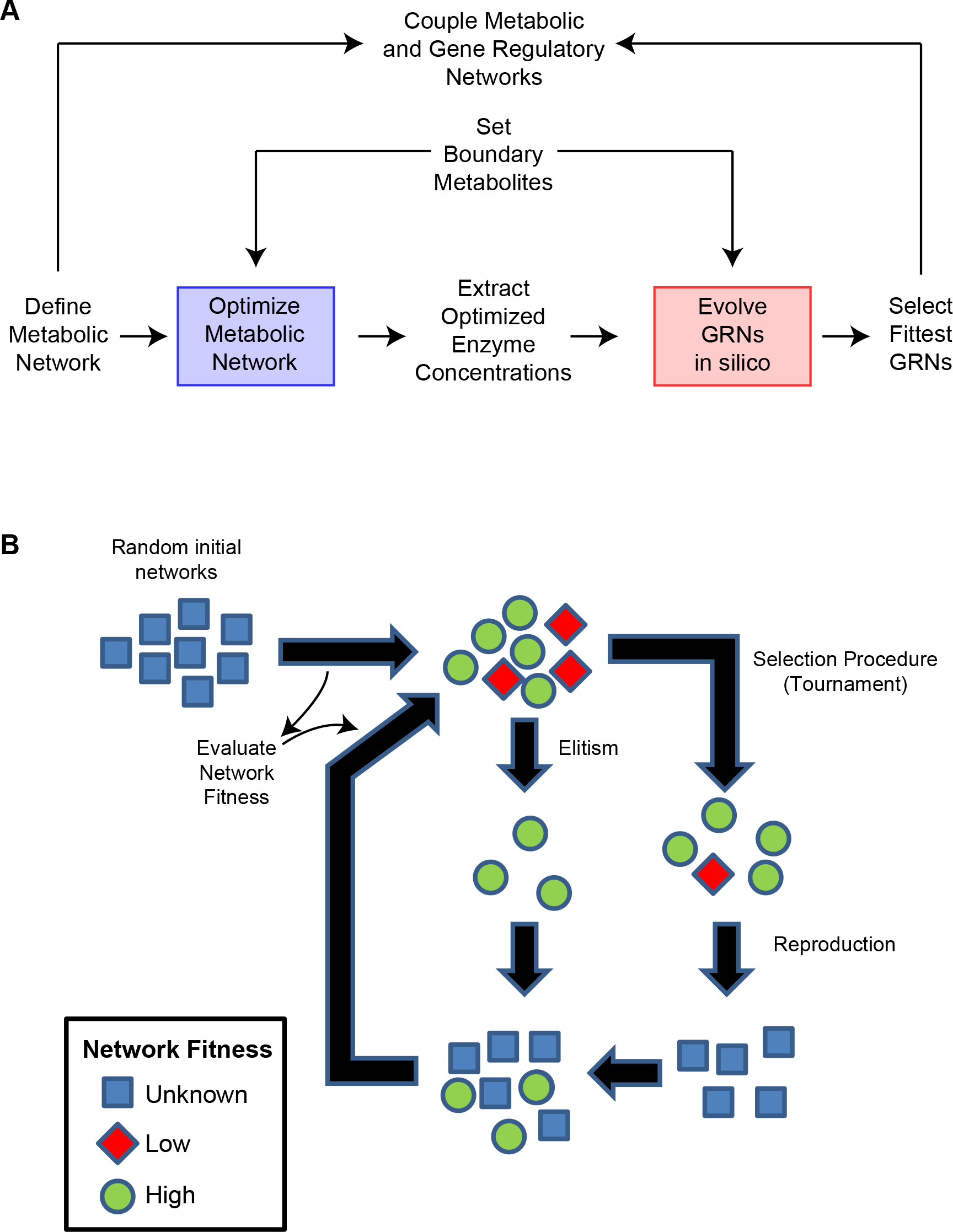
Pipeline of designing self-optimizing integrated GRN-metabolic networks. (a) The two boxes represent the major algorithms used to (1) optimize metabolic fluxes (blue box) and to (2) train GRNs for desired input-output characteristics (red box). (b) Pipeline of in silico evolution of the GRNs.

### Optimization of metabolic fluxes under the total enzyme mass constraint

In our model systems, metabolic reactions were based on Michaelis-Menten and Hill kinetics. The enzymes regulating the metabolic networks were assumed to maintain their total enzyme mass at the level of 10*^−^*^19^ kg per cell, which corresponds to 600 enzymes (under the assumption that the molar mass of all the enzymes is 100 kDa). Under this enzyme mass constraint, we obtained enzyme distributions that maximize metabolic fluxes in linear metabolic networks by using an optimization algorithm based on the Brown’s result, Equation (S14). This optimization procedure was performed for a range of boundary metabolite concentrations to obtain the characteristics between these concentrations and optimal enzyme distributions. For each metabolic network there is only one enzyme distribution where Equation (S14) holds, given a set of kinetic parameters [12]

In the optimization method, (1) scaled flux control coefficients were computed at the steady-state, divided by the total enzyme mass, and subsequently ranked. (2) The enzyme concentration with the highest value was increased and the enzyme concentration with th lowest value was decreased such that the total enzyme mass remained constant. The amount of change in the enzyme concentration was kept sufficiently small (0.1 – 0.5%), since flux control coefficients change with enzyme concentrations. This procedure of steps (1) and (2) was iterated until the newly ranked values were identical within a given convergence error. See SI Section S2 for more details of this algorithm.

### In silico evolution of GRNs

To obtain GRNs that regulate and maintain the metabolic networks at their optimal flux levels, we used the concentrations of the boundary metabolites as input signals of the GRNs and the optimized enzyme distributions as the output signals. These input-output characteristics were applied as a training data set to evolve and select desired GRNs. First, a population of random biochemical network topologies was built. Then, selection and mutation were used to evolve the topologies and perturb reaction parameters, to be trained to follow the input-output characteristics.

Specifically, random GRNs with a population size *N* was built. These GRNs were modeled based on Hill kinetics [27, 33, 34]. Up to three regulators were permitted to control gene expression by performing AND or OR operations [35] with non-competitive (independent) binding. For example, AND operation between two activators exists if both the activators cannot bind individually, but must be dimerized before they can bind to the promoter region. By contrast, OR operation exists when one activator is sufficient for activation, whereas presence of both activators enhances expression.

To evolve GRNs that optimize the distribution of the enzymes regulating the metabolic network, the degree of fitness was quantified by a performance error *F* in the input-output characteristics:

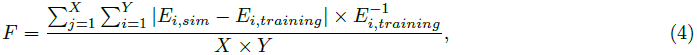

where the absolute difference between the desired and trained levels of each (*i*-th) enzyme is divided by the desired enzyme level, and then summed over all training input-output pairs (the number of the pairs is *X*). Finally, this value is divided by the number of boundary metabolite conditions (*X*) and the number of enzymes (*Y*) to allow comparing models with a different number of boundary metabolite conditions and/or number of enzymes. In other words, the algorithm optimized percentage error per boundary metabolite condition per enzyme.

Our *in silico* approach for GRN evolution is schematically described in Figure 8(b). After evaluating the population fitness, a subset of the GRNs was selected to breed for a new generation. Tournament selection was used to select *S* ‘winning’ GRNs, and elitism was used to guarantee survival of the *N−S* fittest GRNs, for a new population of *N* GRNs. Tournament selection was performed by randomly selecting two GRNs and returning the fittest GRN. After performing tournament selection, the *S* ‘winners’ were subject to structural mutations (insertion, deletion, and transmutation of activators and repressors) and parameter mutations (bounded randomization of kinetic coefficients, as outlined below).

Parameters *V_max_*, *V_deg_*, *K_a_*, and *h* (Hill coefficient) were permitted to be randomized between carefully determined boundaries. For example, a parameter value (*p*_before_) can be multiplied by a random value between zero and two to obtain a new one (*p*_after_). These random values are drawn from a uniform distribution. As a result, the mean values of the parameter before and after the perturbation are identical and the standard deviation of the perturbed parameter value (*p*_after_) is equal to 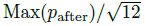. To prevent upwards drift over time a new random value between the minimum and maximum parameter value was assigned if a parameter value mutated to out of boundary values.

### Optimization of fluxes for general metabolic networks

For more general metabolic pathways, a different optimization algorithm was used. In contrast to the previous optimization algorithm in which metabolic flux was optimized under the constraint of fixed total enzyme mass, this optimization algorithm was performed to minimize the total enzyme mass given the constraint of fixed metabolic fluxes. Because these fluxes are fixed in this algorithm, the change in the flux due to parameter perturbations must be zero:

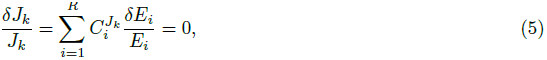

where *k* describes the independent flux and *i* the reaction number. This represents an under-determined system of linear equations, which means that this system cannot be solved for each *δE*. Thus, we semi-randomly selected *V−W* enzymes to perturb their concentrations, where *V* and *W* are the number of enzymes in the system and fixed fluxes, respectively. Then, the remaining enzyme concentrations were computed by using Equation (5). If the total enzyme quantity decreased, we accepted the proposed perturbations. Otherwise, the enzyme concentrations were reset to the previous values. As a consequence, the enzyme concentrations can be updated such that total enzyme mass decreases under the constraint of fixed fluxes.

### Algorithm implementation

The software for performing the optimization algorithms for metabolic networks was developed in Python [36] by using the Python libraries `NumPy` [37], `SymPy` [38], and `PySCeS` [39]. All metabolic and gene regulatory networks were written in the PySCeS Modeling Description Language (MDL). The software for the *in silico* evolution of GRNs was developed in a combination of C++ and Python. This implementation makes use of the Mersenne-Twister pseudo-random number generator [40].

## Acknowledgments

The authors gratefully acknowledge the contributions of NSF (No. 0827592 in Theoretical Biology, No. 1158573 in Molecular and Cellular Biosciences, No. 0527023 in Integrative Biological Research), NWO FOM/ALW 10TBSC24, and the Washington NASA Space Grant. This project was carried out within the research programme of BioSolar Cells, co-financed by the Dutch Ministry of Economic Affairs.

## References

1. Stricker J, Cookson S, Bennett MR, Mather WH, Tsimring LS, et al. (2008) A fast, robust and tunable synthetic gene oscillator. Nature 456: 516–519.

2. Friedland AE, Lu TK, Wang X, Shi D, Church G, et al. (2009) Synthetic gene networks that count. Science 324: 1199–1202.

3. Entus R, Aufderheide B, Sauro H (2007) Design and implementation of three incoherent feed-forward motif based biological concentration sensors. Systems and synthetic biology 1: 119–128.

4. Tabor JJ, Salis HM, Simpson ZB, Chevalier AA, Levskaya A, et al. (2009) A synthetic genetic edge detection program. Cell 137: 1272–1281.

5. Nevozhay D, Adams RM, Murphy KF, Josić K, Balázsi G (2009) Negative autoregulation linearizes the dose–response and suppresses the heterogeneity of gene expression. Proceedings of the National Academy of Sciences 106: 5123–5128.

6. Gardner TS, Cantor CR, Collins JJ (2000) Construction of a genetic toggle switch in escherichia coli. Nature 403: 339–342.

7. Purnick PEM, Weiss R (2009) The second wave of synthetic biology: from modules to systems. Nat Rev Mol Cell Biol 10: 410–422.

8. Sleight S, Bartley B, Lieviant J, Sauro H (2010) Designing and engineering evolutionary robust genetic circuits. Journal of Biological Engineering 4: 12.

9. Ducat DC, Way JC, Silver PA (2011) Engineering cyanobacteria to generate high-value products. Trends in Biotechnology 29: 95–103.

10. Hellingwerf K, Teixeira de Mattos M (2009) Alternative routes to biofuels: Light-driven biofuel formation from co2 and water based on the ‘photanol’ approach. Journal of Biotechnology 142: 87–90.

11. Ludwig M, Bryant DA (2011) Transcription profiling of the model cyanobacterium synechococcus sp. strain pcc 7002 by nextgen (solid) sequencing of cdna. Frontiers in Microbiology 2.

12. Brown GC (1991) Total cell protein concentration as an evolutionary constraint on the metabolic control distribution in cells. Journal of Theoretical Biology 153: 195–203.

13. E Klipp, Heinrich R (1999) Competition of enzymes in metabolic pathways: Implications for optimal distributions of enzyme concentrations and for the distribution of flux control. BioSystems 54: 1–14.

14. Beg Q, Vazquez A, Ernst J, De Menezes M, Bar-Joseph Z, et al. (2007) Intracellular crowding defines the mode and sequence of substrate uptake by escherichia coli and constrains its metabolic activity. Proceedings of the National Academy of Sciences 104: 12663–12668.

15. Atkinson DE, Roach PJ, Schwedes JS (1975) Metabolite concentrations and concentration ratios in metabolic regulation. Advances in Enzyme Regulation 13: 391–411.

16. Schuster S, Schuster R, Heinrich R (1991-04-01) Minimization of intermediate concentrations as a suggested optimality principle for biochemical networks. Journal of Mathematical Biology 29: 443–455.

17. Heinrich R, Holzhuütter HG, Schuster S (1987) A theoretical approach to the evolution and structural design of enzymatic networks; linear enzymatic chains, branched pathways and glycolysis of erythrocytes. Bulletin of Mathematical Biology 49: 539–595.

18. Heinrich R, Klipp E (1996) Control analysis of unbranched enzymatic chains in states of maximal activity. Journal of Theoretical Biology 182: 243–252.

19. Eldar A, Elowitz M (2010) Functional roles for noise in genetic circuits. Nature 467: 167–173.

20. Fell D (1992) Metabolic control analysis: a survey of its theoretical and experimental development. Biochem J: 313–330.

21. Kacser H, Burns J (1995). The control of flux.

22. Fell D, Thomas S (1995) Physiological control of metabolic flux: the requirement for multisite modulation. Biochem J: 35–39.

23. Moreno-Sánchez R, Saavedra E, Rodrguez-Enríquez S, Gallardo-Pérez J, Quezada H, et al. (2010) Metabolic control analysis indicates a change of strategy in the treatment of cancer. Mitochondrion 10: 626–639.

24. Kim K, Sauro H (2012) Adjusting phenotypes by noise control. PLoS computational biology 8: e1002344.

25. Kim K, Sauro H (2010) Sensitivity summation theorems for stochastic biochemical reaction systems. Mathematical biosciences 226: 109–119.

26. Deckard A, Sauro HM (2004) Preliminary studies on the in silico evolution of biochemical networks. ChemBioChem 5: 1423–1431.

27. Paladugu SR, Chickarmane V, Deckard A, Frumkin JP, McCormack M, et al. (2006) In silico evolution of functional modules in biochemical networks. Systems biology 153: 223–235.

28. Shen-Orr SS, Milo R, Mangan S, Alon U (2002) Network motifs in the transcriptional regulation network of escherichia coli. Nat Genet 31: 64–68.

29. Ronen M, Rosenberg R, Shraiman BI, Alon U (2002) Assigning numbers to the arrows: Parameterizing a gene regulation network by using accurate expression kinetics. Proceedings of the National Academy of Sciences 99: 10555–10560.

30. Zaslaver A, Mayo AE, Rosenberg R, Bashkin P, Sberro H, et al. (2004) Just-in-time transcription program in metabolic pathways. Nat Genet 36: 486–491.

31. Babu M (2008) Evolutionary and temporal dynamics of transcriptional regulatory networks. Bio-Inspired Computing and Communication: 174–183.

32. Goldberg D (1989) Genetic algorithms in search, optimization, and machine learning. Addison-wesley.

33. Rosenfeld N, Elowitz M, Alon U (2002) Negative auto-regulation speeds the response times of transcription networks. J Mol Biol 323: 785–793.

34. Gjuvsland A, Hayes B, Omholt S, Carlborg O (2007) Statistical epistasis is a generic feature of gene regulatory networks. Genetics 107: 411–420.

35. Sauro H (2011) Enzyme Kinetics for Systems Biology. analogmachine.org.

36. van Rossum G, et al. (1991). The Python Language Reference. URL http://www.python.org/.

37. Oliphant TE (2006) Guide to numpy. Technical report, http://www.numpy.org.

38. Certik O (2008) Sympy python library for symbolic mathematics. technical report. Technical report.

39. Olivier B, Rohwer J, Hofmeyr J (2005) Modelling cellcular systems with PySCeS. Bioinformatics 21: 560–561.

40. Matsumoto M, Nishimu T (1998) Mersenne twister: A 623-dimensionally equidistributed uniform pseudo-random number generator. ACM Transactions on Modeling and Computer Simulation: 3–30.

